# Antibiotic concentrations in the sinonasal secretions and tissue in CRS patients after oral therapy: a randomized trial

**DOI:** 10.1101/2020.06.17.158535

**Authors:** Joey Siu, Lilian Klingler, Yi Wang, Cheung-Tak Hung, Soo Hee Jeong, Susan Smith, Malcolm Tingle, Brett Wagner Mackenzie, Kristi Biswas, Richard Douglas

**Affiliations:** Department of Surgery, The University of Auckland, Auckland, New Zealand; Research and Development, Zenith Technology Corporation Limited, Dunedin, New Zealand; Department of Pharmacology and Clinical Pharmacology, The University of Auckland, Auckland, New Zealand; Labtests, Auckland, New Zealand

**Keywords:** Sinusitis, Bacteria, Microbiota, Antibiotics, Antibiotic resistance, Macrolides, Tetracyclines

## Abstract

**Background:** Despite the widespread prescription of antibiotics for patients with chronic rhinosinusitis (CRS), the extent to which drug distribution to the sinonasal mucosa influences their efficacy remains largely undefined.

**Methods:** Thirty subjects undergoing functional endoscopic sinus surgery (FESS) for bilateral CRS were randomized to one of three groups: 1) doxycycline (100 mg daily for seven days) 2) roxithromycin (300 mg daily for seven days) and 3) control (no antibiotics given). Drug levels were measured using liquid chromatography-tandem mass spectrometry in sinonasal secretions, sinonasal tissues and serum at steady state. Nasal endoscopy (Modified Lund-Kennedy) and Gastrointestinal Symptom Rating Scale (GSRS) scores were recorded.

**Results:** Antibiotic concentrations in the nasal secretions were significantly lower compared to those in the serum and tissue (mean mucus/serum ratio at steady state = 0.16 and 0.37 for doxycycline and roxithromycin respectively; *p*<0.01). A short course of antibiotic intake did not correlate with any difference in clinical outcomes except where slightly higher GSRS scores were reported in the roxithromycin group (*p*=0.04).

**Conclusions:** Although the efficacy of doxycycline and roxithromycin in sinonasal mucus *in vivo* cannot be predicted solely from reported minimum inhibitory concentrations, given the added complication of bacterial biofilm antimicrobial tolerance, these results suggest that low mucosal penetration of antibiotics may be one of the factors contributing to the limited efficacy of these agents in the treatment of CRS.

## INTRODUCTION

Despite the widespread use of antibiotics in the treatment of chronic rhinosinusitis (CRS), their efficacy for this indication remains debatable. There is increasing evidence that the repeated use of broad-spectrum antibiotics is associated with both microbial dysbiosis and the emergence of resistant bacterial strains(1-6). The changes that occur in the sinonasal microbiota during oral antibiotic treatment in CRS patients are poorly understood, and the microbiological effect of antibiotics at a molecular level have not been correlated with clinical outcome measures.

Chronic rhinosinusitis (CRS) represents a spectrum of disorders that result from a variety of immunopathological mechanisms that lead to persistent inflammation of the paranasal sinus mucosa. Its clinical classification in two broad groups: CRS with nasal polyps (CRSwNP) and CRS without NPs (CRSsNP) provides insight into the severity of disease, extensiveness of surgery required, and the efficacy of medical therapies. On a cellular level, both of these phenotypes are characterized by epithelial disruption, ciliary dysfunction, mucus gland hyperplasia, bacterial overgrowth and the formation of biofilms(7). The role of microbes in the pathogenesis of CRS is largely unknown, but bacteria probably contribute to the persistence and severity of the disease(8). The International Consensus Statement on Allergy and Rhinology recommends the use of oral macrolides as an option in the treatment of CRS without nasal polyps (CRSsNP) on the basis that they have shown at least temporary benefit in some studies, reducing endoscopy scores and improving symptoms(9). A reduction in polyp size with doxycycline has been demonstrated in patients with polyps (CRSwNP), but no difference was found in patient-reported outcomes(10). The benefit of a short-term period (<3 weeks) of oral antibiotics is particularly unclear(9). Several studies report that a short-term use of antibiotics from various antibiotic classes improve clinical symptoms such as nasal discharge and nasal blockage but there is a lack of clinical trials showing a direct benefit in improving the patient’s intraoperative condition(11-13).

The extent to which drug distribution to the sinonasal mucosa influences the efficacy of oral antibiotics in patients with CRS remains largely undefined. In vitro bacterial susceptibility testing does not take into account the pharmacokinetics of the antimicrobial agent and the variability in drug distribution to various sites in the body(14). Current analytical methodologies and instrumentation allow accurate quantitative analysis of drug concentrations in nasal secretions and tissue by chromatography and spectrometry(15-30). However, few studies have used such methods to determine sinonasal concentrations of antibiotics used in CRS, particularly those from the macrolide and tetracycline groups(26, 31, 32). Although the studies performed to date have suggested a therapeutic concentration of antibiotics greater than reported minimum inhibitory concentrations (MIC) was achieved, there is little evidence supporting their efficacy *in vivo*.

This randomized control trial aimed to determine the concentrations of two commonly prescribed antibiotics, doxycycline and roxithromycin, in the nasal secretions, serum and sinonasal tissues of CRS patients following a one-week course. The clinical impact of a short-term duration of antibiotics on nasal endoscopy scores (Modified Lund-Kennedy), and adverse gastrointestinal symptoms as measured by the Gastrointestinal Symptom Rating Scale (GSRS) were secondary endpoints. Finally, further analysis was performed on a subset of five patients, initiating an investigation into the relationship between drug concentrations and patient-specific antibiotic susceptibility from measured MIC’s of predominating sinonasal bacteria. The data presented in the sub analysis are part of an ongoing study aiming to test our hypothesis that MIC’s may have limited relevance to CRS patients in the setting of biofilms.

## MATERIALS AND METHODS

### Study design and sample collection

Thirty subjects undergoing functional endoscopic sinus surgery (FESS) for extensive bilateral CRS were recruited for this study (Table 1). This clinical population was deemed most suitable for the measurement of antibiotic concentrations in various tissue sites which could be sampled at the time of their operation.

**Table 1:**
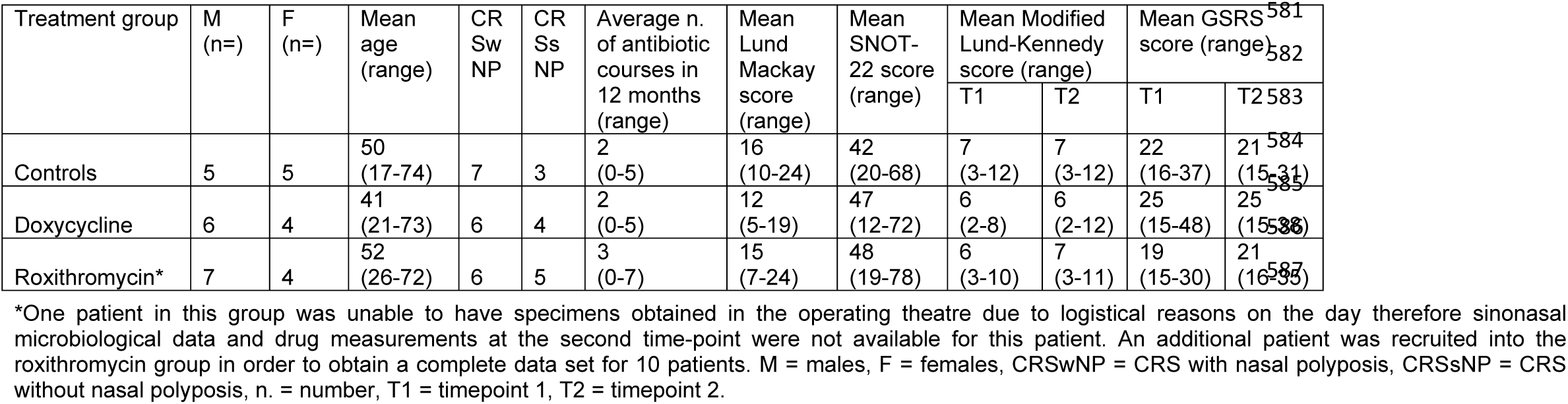
Demographics, disease state and average clinical scores of study cohort.

| Treatment group | M<br>(n=) | F<br>(n=) | Mean<br>age<br>(range) | CR<br>Sw<br>NP | CR<br>Ss<br>NP | Average n.<br>of antibiotic<br>courses in<br>12 months<br>(range) | Mean<br>Lund<br>Mackay<br>score<br>(range) | Mean<br>SNOT-22 score<br>(range) | Mean Modified<br>Lund-Kennedy<br>score (range) |  | Mean GSRS<br>score (range) |  |
| --- | --- | --- | --- | --- | --- | --- | --- | --- | --- | --- | --- | --- |
|  |  |  |  |  |  |  |  |  | T1 | T2 | T1 | T2 |
| Controls | 5 | 5 | 50<br>(17-74) | 7 | 3 | 2<br>(0-5) | 16<br>(10-24) | 42<br>(20-68) | 7<br>(3-12) | 7<br>(3-12) | 22<br>(16-37) | 21<br>(15-31) |
| Doxycycline | 6 | 4 | 41<br>(21-73) | 6 | 4 | 2<br>(0-5) | 12<br>(5-19) | 47<br>(12-72) | 6<br>(2-8) | 6<br>(2-12) | 25<br>(15-48) | 25<br>(15-38) |
| Roxithromycin* | 7 | 4 | 52<br>(26-72) | 6 | 5 | 3<br>(0-7) | 15<br>(7-24) | 48<br>(19-78) | 6<br>(3-10) | 7<br>(3-11) | 19<br>(15-30) | 21<br>(16-35) |

\*One patient in this group was unable to have specimens obtained in the operating theatre due to logistical reasons on the day therefore sinonasal microbiological data and drug measurements at the second time-point were not available for this patient. An additional patient was recruited into the roxithromycin group in order to obtain a complete data set for 10 patients. M = males, F = females, CRSwNP = CRS with nasal polyposis, CRSsNP = CRS without nasal polyposis, n. = number, T1 = timepoint 1, T2 = timepoint 2.

Patients aged <16 years, with acute exacerbations, smokers, or who had been prescribed oral antibiotics or systemic corticosteroids during the four weeks prior to recruitment were excluded. Patients with a diagnosis of cystic fibrosis, fungal sinusitis, chronic kidney disease, impaired liver function, immunodeficiency, congenital mucociliary problems, systemic vasculitis and granulomatous disease, chronic gastrointestinal inflammatory or immune-mediated diseases were also excluded. This study was approved by the New Zealand Health and Disability Ethics Committee (17/NTB/228) and written informed consent was obtained from all participants and in the case of a single participant aged <18 (16 years) in the company of his parents.

Eligible patients were randomised using a random number generator to one of three groups: 1) doxycycline (100 mg orally with food daily for seven days) 2) roxithromycin (300 mg orally at least 30 minutes before food daily for seven days) and 3) control (no treatment). Patients were diagnosed according to the 2012 European Position Paper (EPOS) definition of CRS(33). Patients in the medication groups started their medications seven days before surgery, taking one dose every morning up to and including the morning of surgery. All patients were recruited during a pre-operative consultation fewer than four weeks before FESS surgery (timepoint 1). During FESS, multiple specimens were collected from the patients in the medication groups (timepoint 2). Samples collected include blood, bilateral middle meatal mucus samples, inferior turbinates and ethmoid bulla mucosa. Patients completed a gastrointestinal symptom rating scale (GSRS)(34), and were assessed endoscopically at both timepoints 1 and 2. The GSRS is a 15-item questionnaire for patients with gastrointestinal symptoms validated for various gastrointestinal disorders(34). The sum of the scores ranging from 15 to 105 is regarded as the total score. Nasal endoscopy scores are derived from the Modified Lund-Kennedy System (MLK) for polyps, discharge, and oedema on a scale of 0– 12(35). Baseline rhinosinusitis symptoms(35) (SNOT-22), radiological Lund-Mackay scores(36) and patient demographic data were collected (Table 1).

One patient underwent limited FESS (bilateral uncinectomy, middle meatal antrostomy, anterior ethmoidectomy) and the remaining underwent comprehensive FESS including ethmoidectomy, sphenoidectomy, and frontal sinus dissection. Four of these were revision operations and five of them included frontal sinus drill out surgery. All study investigators apart from those conducting drug assays of biological specimens were kept blinded to randomization until completion of the study. Those conducting drug assays required knowledge of the drug since the assays were drug specific. To maximize patient compliance, it was elected not to blind patients to their treatment group. The benefit of this in optimizing the determination of drug concentrations outweighed the potential impact on clinical correlation with the GSRS score, considered one of the secondary endpoints in the study. Patients were asked to keep their medication packets which were checked for compliance during the consultation at timepoint 2. In accordance with standard treatment for CRS, all patients continued to use daily topical corticosteroid nasal sprays and performed regular sinonasal saline lavage, excluding the day of FESS surgery.

Samples were collected in the operating room prior to the application of topical vasoconstrictors and anaesthetic solutions. Undiluted nasal secretions from both sides of the sinonasal cavity were obtained by aspiration and collected in a mucus trap extractor. The extractor was weighed before and after the collection of secretions in order to calculate the weight of sample collected for each patient. Tissue samples (ethmoid bulla and inferior turbinates) were collected bilaterally using standard surgical techniques as part of the standard FESS procedure. Blood was withdrawn using a 4 mL serum tube without clot activator. All sinonasal specimens and blood specimens were transported on ice to the laboratory within 2 h. Blood was centrifuged in order to retain the serum component for storage. All specimens were stored at -80 °C until drug assay.

### Determination of antibiotic concentrations by LC-MS/MS

A sensitive liquid chromatography-tandem mass spectrometry (LC-MS/MS) assay was employed to determine antibiotic concentrations in human nasal mucus, sinonasal tissue and serum. Samples were separated with a Luna C18 column (Phenomenex, CA, USA) and antibiotic levels were detected with a triple-quadrupole mass spectrometer using electrospray ionization in positive mode and multiple reaction monitoring. QTRAP® 6500 and API 4000™ systems (SCIEX, MA, USA) were employed for doxycycline and roxithromycin respectively. Methacycline was used as the internal standard for doxycycline and clarithromycin was used as the internal standard for roxithromycin. Calibration curves were obtained from partial method validation (lower limit of quantification (LLOQ) for doxycycline = 32ng/mL in serum; 1ng/mL in mucus and tissue, LLOQ for roxithromycin = 0.2µg/mL in serum; 5ng/mL in mucus and tissue). The determination of both drugs in these biological matrices was reliable and reproducible according to pharmaceutical guidelines(15, 37, 38).

#### Sample preparation

Serum sample preparation for the determination of doxycycline concentrations involved deproteinization of serum sample with ACN, drying the sample under a stream of nitrogen before reconstitution of the sample with ACN:B.P. water (15:85, v/v) containing 0.1% (v/v) formic acid. Serum sample preparation for the determination of roxithromycin concentrations involved deproteinization with acetonitrile (ACN) then further dilution of the sample with ACN:B.P. water (30:70, v/v) containing 0.1% (v/v) formic acid. An additional step of dilution with methanol:B.P. water (50:50, v/v) was performed initially for nasal mucus samples, while dilution and homogenization in ACN:B.P. water (50:50, v/v) containing 0.05% of formic acid was performed initially for the sinonasal tissue samples.

### Subgroup analysis of bacterial isolates and their antibiotic susceptibilities

A random block of five patients taking either doxycycline or roxithromycin had further samples taken for bacterial culture analysis and antibiotic susceptibility testing. This included a right-sided middle meatus swab at timepoints 1 and 2 as well as a right sided ethmoid bulla tissue sample at timepoint 2. The presence or absence of *Staphylococcus aureus* and beta-haemolytic *Streptococci* on these samples were reported using colonial appearances and confirmed using Matrix-Assisted Laser Desorption/Ionization-Time of Flight (MALDI-TOF). All other organisms were only isolated for genus identification using MALDI-TOF if they were considered a dominant organism. A dominant organism grew two times more than the other organisms present. Growths were defined as light (growth only in the initial inoculum), moderate (growth in the initial inoculum and streak lines but not over the entire plate) or heavy (growth over entire plate). The minimum inhibitory concentration required to inhibit the growth of 90% of organisms (MIC^90^) of all identified organisms to doxycycline, erythromycin and amoxicillin-clavulanic acid were tested using MIC strips (Liofilchem®) according to manufacturer instructions. Bacterial susceptibilities were interpreted using the European Committee on Antimicrobial Susceptibility Testing (EUCAST) criteria(39). Erythromycin was selected over roxithromycin as the closest macrolide drug due to the unavailability of roxithromycin test strips.

### Statistical analyses

Significance levels were set to *p*<0.05 (two-sided). Patient data were summarised descriptively, for continuous and categorical variables. The student’s *t* test was applied to analyse differences in continuous variables between groups. The chi-squared test was similarly used for categorical variables. Linear regression analyses were conducted to determine correlations between drug concentrations and clinical scores (MLK and GSRS scores) in the medication groups. Based on an *α* value of .05, a sample size estimate indicated that a correlation coefficient of 0.77 could be detected with a sample size of 10 at a power of 0.80.

## RESULTS

All patients complied with trial protocol and completed the study. There were no statistically significant differences in gender, age, disease type (CRSwNP vs. CRSsNP), presence of asthma, baseline clinical scores (Lund-Mackay, MLK, SNOT-22 and GSRS) or number of antibiotic courses taken in the 12 months preceding recruitment between all three patient groups. The roxithromycin group had a slightly worse GSRS score change compared with the control group (2.3 vs. -1.9; *p*=0.04). No other significant differences were observed within or between study groups for changes in clinical scores (MLK and GSRS).

Regression analyses were performed comparing clinical score changes (MLK, and GSRS) with antibiotic concentrations in the nasal mucus, tissue and blood. These analyses demonstrated a reduction in the GSRS score (R^2^ = 0.45, *p*=0.04) with increasing mucus antibiotic concentrations in the doxycycline group (Figure 1). No other significant correlations were found.

**Figure 1.**
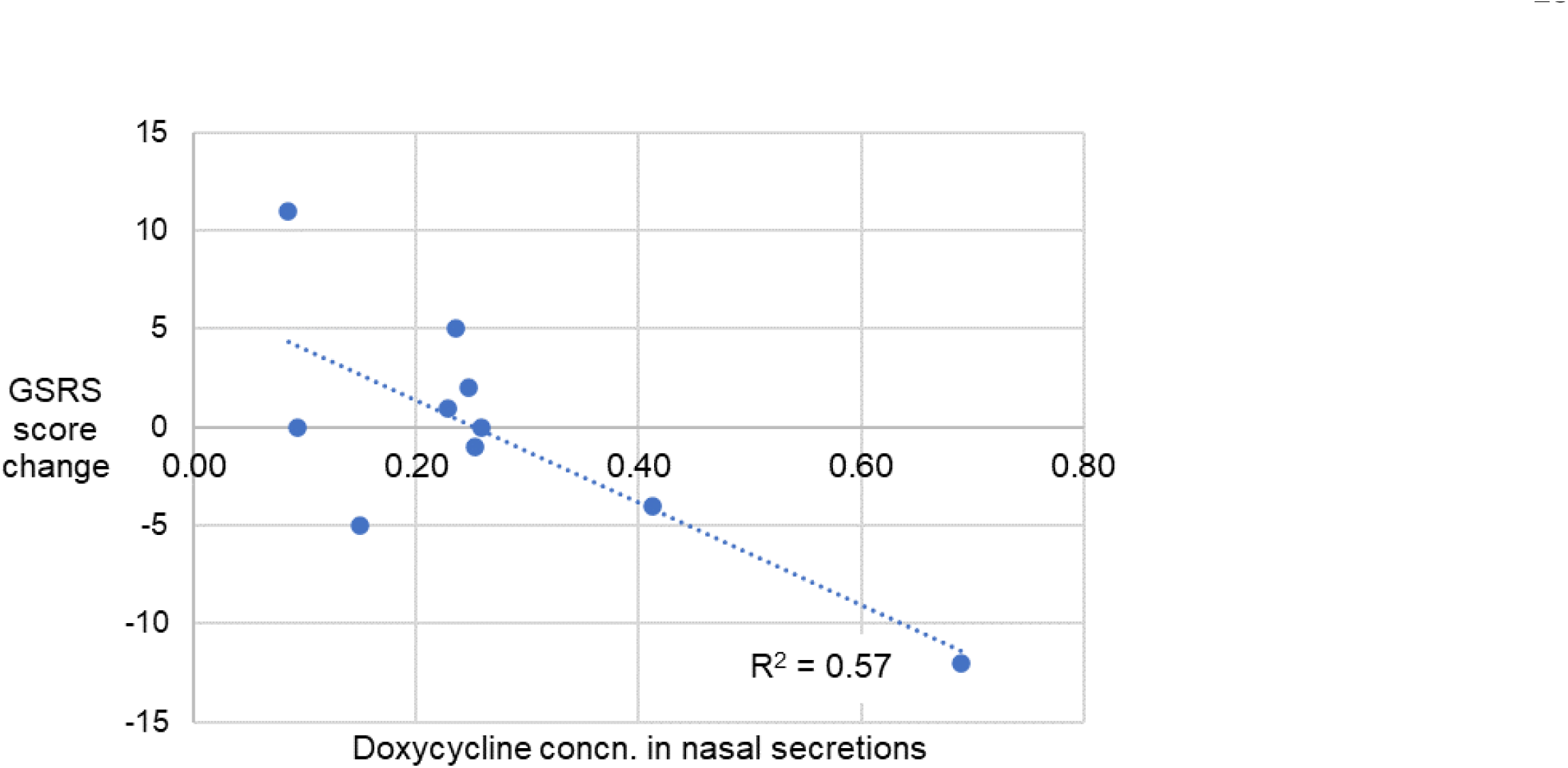
Gastrointestinal symptoms rating scale (GSRS) score change vs. doxycycline concentrations in nasal secretions.

### Drug penetration in nasal secretions and sinonasal tissue

Mean antibiotic concentrations and penetration are summarized in Table 2 and Figure 2. Individual data are included in Supplementary Table S1 (for reviewers’ information only). The median time of specimen sampling from the last dose of medication was 6.5 h (range, 2-15). There was no correlation between time of sampling to drug concentration in the serum, mucus or sinonasal tissue specimens. The mean concentrations (± SD) of doxycycline detected were – serum: 1.6 ± 0.9 µg/mL (range, 0.5-3.0); mucus: 0.27 ± 0.18 µg/mL (range, 0.09-0.69); turbinates: 1.4 ± 0.48 µg/mL (range, 0.49-2.1); ethmoid bullae: 1.6 ± 0.5 µg/mL (range, 0.7-2.4). The mean concentrations (±SD) of roxithromycin detected were – serum: 4.3 ± 1.1 µg/mL (range, 2.0-5.5); mucus: 1.6 ± 1.7 µg/mL (range, 0.1-4.8); turbinates: 2.8 ± 0.9 µg/mL (range, 0.9-4.3); ethmoid bullae: 2.6 ± 1.1 µg/mL (range, 0.7-4.8).

**Table 2.** Mean antibiotic concentrations and penetration ratios.

|  | Doxycycline | *p* value | Roxithromycin | *p* value |
| --- | --- | --- | --- | --- |
| Time of sampling (hrs since last dose) ± SD | 8 ± 3<br>(range, 6-15<sup>a</sup>) | | 6 ± 3<br>(range, 2-9) | |
| Serum concentration (μg/mL) ± SD | 1.62 ± 0.91<br>(range, 0.46-2.99) | | 4.28 ± 1.12<br>(range, 2.07-5.52) | |
| Mucus concentration (μg/mL) ± SD | 0.27 ± 0.18<br>(range, 0.09-0.69) | | 1.60 ± 1.67<br>(range, 0.08-4.76) | |
| Turbinate tissue concentration (μg/mL) ± SD | 1.38 ± 0.48<br>(range, 0.49-2.05) | | 2.76 ± 0.90<br>(range, 0.85-4.29) | |
| Ethmoid bulla tissue concentration (μg/mL) ± SD | 1.59 ± 0.54<br>(range, 0.68-2.42) | | 2.62 ± 1.13<br>(range, 0.72-4.83) | |
| Mucus-to-serum ratio | 0.16<br>(range, 0.08-0.67) | <0.001 | 0.37<br>(range, 0.02-1.06) | 0.002 |
| Tissue-to-mucus ratio | 5.58<br>(range, 3.09-7.52) | <0.0001 | 1.68<br>(range, 0.66-29.7) | <0.001 |
| Tissue-to-serum ratio | 0.91<br>(range, 0.56-1.61) | 0.2 | 0.63<br>(range, 0.37-1.08) | <0.001 |

<sup>a</sup>This patient had a much longer sampling time since their operation was scheduled for the morning and they were instructed to take the last dose of doxycycline with their meal the night before surgery.

**Figure 2.**
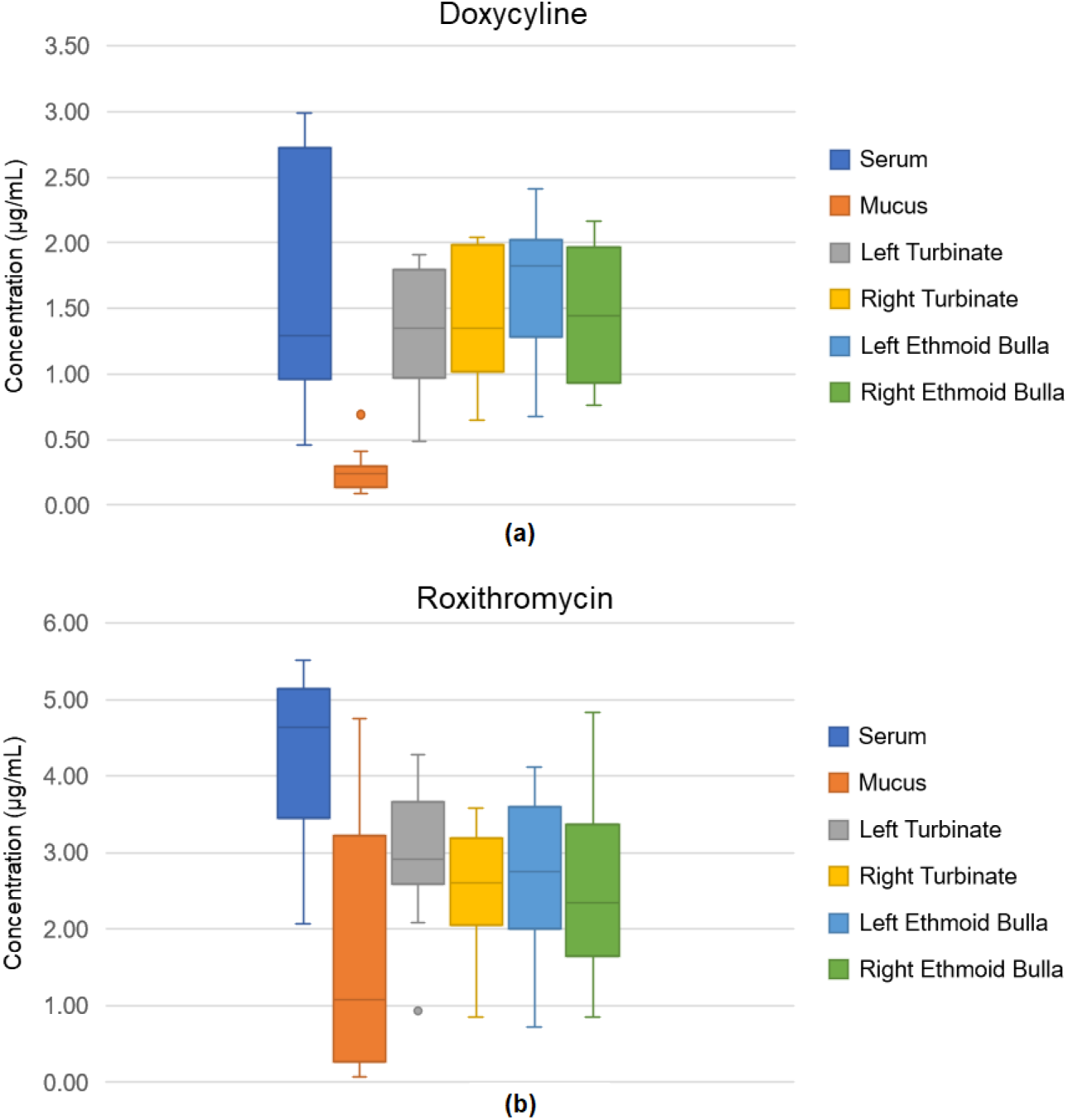
Antibiotic concentrations in the serum, mucus and different sinonasal tissue sites in the doxycycline (a) and roxithromycin (b) groups.

In the doxycycline group, the mean mucus-to-serum ratio was 0.2 (range, 0.08-0.7). This difference was highly statistically significant (*p*<0.001). The mean tissue-to-mucus ratio was 5.6 (range, 3.1-7.5) while the mean tissue-to-serum ratio was 0.9 (range, 0.6-1.6). There were significantly higher levels of drug measured in the tissue compared to mucus (*p*<0.0001) but not compared to the serum (*p*=0.2). In the roxithromycin group, the mean mucus-to-serum ratio was 0.4 (range, 0.02-1.1). This difference was statistically significant (*p*=0.002). The mean tissue-to-mucus ratio was 1.7 (range, 0.7-30) while the mean tissue-to-serum ratio was 0.6 (range, 0.4-1.1). These differences were highly statistically significant (*p*<0.001).

There was no significant difference in drug concentrations between left and right sinonasal tissue specimens nor between tissue sites (ethmoid bulla versus inferior turbinate).

### Antibiotic susceptibilities of bacterial isolates

A subgroup analysis of bacterial isolates was performed in five randomly selected patients assigned to an antibiotic treatment group (Table 3). A light growth of *S. aureus* was identified in baseline swabs for two patients, which were sensitive to both doxycycline and roxithromycin. These patients (patients 19 and 27) had therapeutic concentrations of antibiotics in the mucus and tissue according to the susceptibility results and *S. aureus* was not detected in any of their post-drug samples. Two patients in the doxycycline group (patients 19 and 23) had bacteria predominating the culture plates for their post-drug samples despite one week of medication. These were moderate growths of *Propionibacterium* and *Moraxella* species respectively and were present despite both being sensitive to doxycycline. Serum, mucus and tissue doxycycline concentrations for the two patients were at least two times greater than the MIC thresholds of the respective organisms.

**Table 3.**
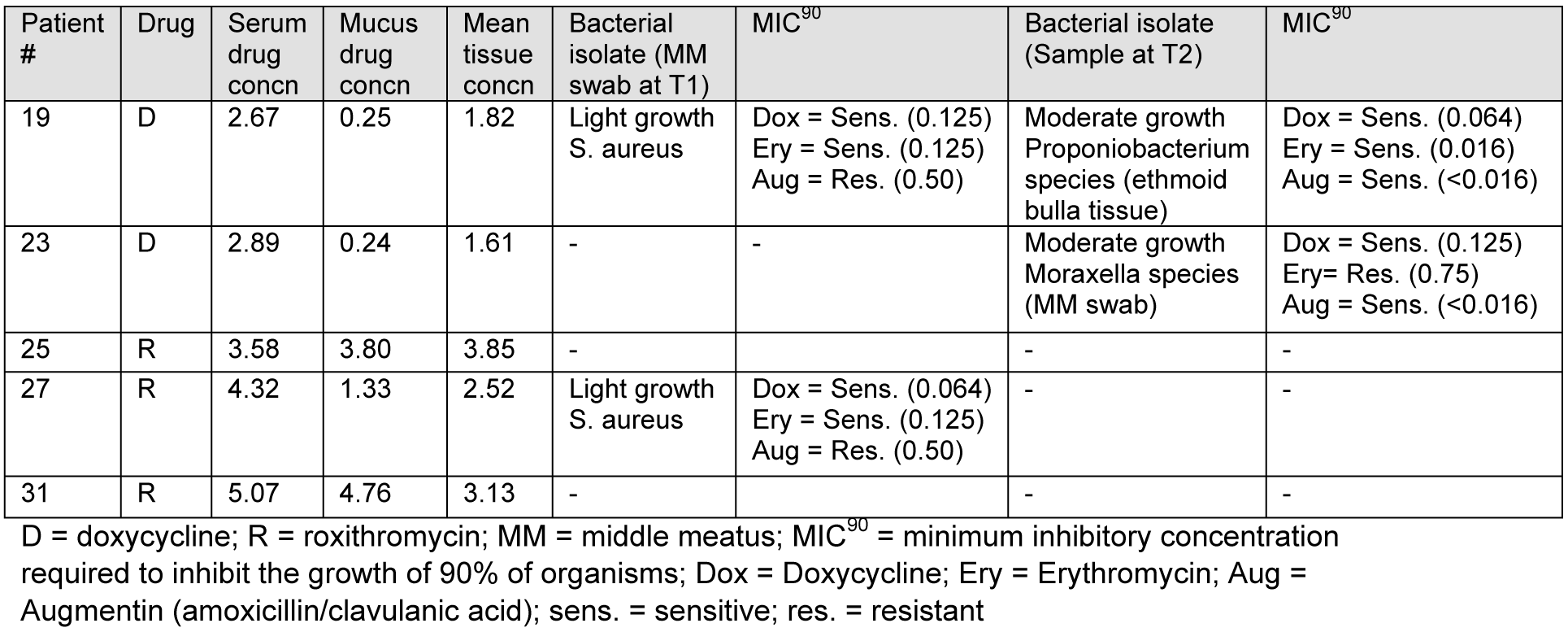
Bacterial isolates cultured from random subgroup and respective bacterial susceptibilities to doxycycline, erythromycin and amoxicillin/clavulanic acid.

## DISCUSSION

The results of this study suggest that the levels of doxycycline and roxithromycin in the mucus of patients with CRS is substantially lower than the level measured simultaneously in the sinonasal mucosa and serum. This finding may provide part of the explanation why antibiotics have not proven to be particularly effective in the treatment of CRS.

Measuring antibiotic concentrations from paranasal tissue and secretions is challenging due to difficulties in various stages of bioanalysis related to small volume samples and the heterogeneous, protein-rich nature of the samples. Few studies have examined antibiotic concentrations in human sinonasal tissues and/or mucus(18-32, 40-44). The majority of these studies were performed in patients with acute sinusitis, acute exacerbations of CRS or upper respiratory tract infections. As a result, data for various antibiotics commonly used for the treatment of CRS are lacking, particularly those from the macrolide and tetracycline groups, compared with fluoroquinolone and beta-lactam drugs(26, 31, 41, 44).

The results from this study showed that doxycycline and roxithromycin concentrations in the nasal secretions were significantly lower compared to that in the serum. The mean ratios at steady state were 0.16 and 0.37 respectively. These values are comparable to those found in bronchial secretions in lung studies(45-48). However, much higher penetration in human nasal secretions are reported for other antibiotics that have been studied(18-32, 40-44). Notably, sinonasal secretion to blood ratios of other macrolides such as telithromycin, clarithromycin and azithromycin have been reported as ≥ 1.0(26, 31). Our results suggest that penetration of doxycycline and roxithromycin in sinonasal tissues were much higher compared to that in the nasal secretions, although tissue levels were still ≤ 1.0 compared to serum, contrary to the published existing studies of several different antibiotics(19, 21, 22, 24, 26, 30). In one study of telithromycin, the ratio of the tissue versus plasma for area under the curve (AUC) was 5.9 for nasal mucosa and 1.6 for ethmoid bone(26). Regardless of drug class, different antibiotics display high variability in tissue site versus intravascular ratios. Nevertheless, there are limitations in the interpretation and comparison of existing studies including small sample sizes, heterogeneity of study populations, specific antibiotic and dosage regimens, nasal sampling methods, time of sampling in relation to the time of dose, pharmacodynamic outcomes and methods of drug analysis.

The analysis of drugs or their metabolites in extravascular compartments such as the mucus and mucosa can improve our understanding of penetration and likely efficacy at the site of disease(17). This statement has been supported by existing studies of pharmacodynamics and pharmacokinetics of antibiotics used to treat respiratory infectious diseases(45-50). It is unknown whether the ability of an antibiotic to kill bacteria above MIC thresholds is more important at the tissue or mucus level within the sinonasal cavity, however existing respiratory studies conclude that epithelial lining fluid is the site where pathogenic bacteria reside(45-50). Given that antibiotic intervention has not been proven to be generally effective in CRS and our results showing that concentrations in the sinonasal mucus were significantly lower compared to those in the serum and tissue we hypothesize that the mucus may be a more important target compartment for antibiotics. This is further supported by our findings suggesting that pathogens commonly associated with acute sinusitis and upper respiratory infection, such as *Staphylococcus aureus, Streptococcus pneumoniae, Streptococcus pyogenes, Haemophilus influenzae* and *Moraxella catarrhalis*, are susceptible to tissue drug levels in relation to MIC^90^ levels for susceptible strains reported in the literature(51-54). On the other hand, for the majority of patients the concentrations we found in mucus was less than the MIC of these species, with the exception of *Streptococcus pneumoniae* and *Streptococcus pyogenes* in the roxithromycin group(51-54). Roxithromycin also exceeded the MIC in mucus of *Corynebacterium spp*., *Lactobacillus spp*. and *Propionibacterium acnes*(51), the presence of which may be important in maintaining a stable sinonasal bacterial community in CRS patients(55). The implications for diversity depletion and microbial dysbiosis should be considered when prescribing these broad spectrum antibiotics(1-6). Additionally, interpretation of the MIC may be misleading as many of the bacteria in the sinonasal mucosa may be living in biofilms and have a substantially higher resistance to antibiotics than the in vitro MIC against planktonic organisms would suggest(7). This is supported by our small subgroup analysis of bacterial isolates in which two patients taking doxycycline were found to have moderate dominant growths of certain organisms despite susceptibility testing showing the MIC for doxycycline was exceeded. In addition, serum, mucus and tissue antibiotic concentrations for these patients were at least two times greater the MIC^90^ thresholds of the respective organisms.

In this study, a positive linear correlation was found between doxycycline concentrations in the nasal mucus and score reduction in the GSRS questionnaires. This suggests a reduction in gastrointestinal effects with increasing penetration of doxycycline into the mucus. Further study is required to investigate the possibility that an increased distribution of antibiotics to the mucus is accompanied by a reduction in the distribution of antibiotics to the gastrointestinal tract. Antibiotic intake did not correlate with any significant difference in objective clinical assessment by nasal endoscopy nor gastrointestinal symptom scores between groups overall except where slightly less favourable gastrointestinal scores were found in patients taking roxithromycin. A larger double-blinded study with a longer medication period is required to validate these clinical correlations, and any differences among patient subgroups such CRSwNP compared to CRSsNP, where the penetration of drugs into polypoid tissue compared with non-polypoid tissue would be of additional interest.

## Limitations

There are some limitations associated with this study. Since patients were not blinded, the interpretation of patient reported GSRS scores is limited. However, the primary outcome of the study was to investigate drug distribution and whether tissue or mucus levels reached reported MICs’ of the antibiotic for common pathogens associated with CRS, while correlation with clinical scores (MLK and GSRS) was considered a secondary endpoint. In addition, major clinical changes were not expected with a brief course of antibiotics. A further secondary analysis was performed on a subset of five patients adding objective data from bacterial swabs. This aimed to provide an insight into the relationships between drug concentrations, patient-specific MIC’s, and bacterial eradication. A follow-up study is required to establish these relationships, since our preliminary results support the hypothesis that MIC’s may have limited relevance to CRS patients in the setting of biofilms

Since samples for drug assays were obtained at the time of surgery, each subject was only available for one sample collection. To best overcome changing penetration ratios over time, sampling was performed at steady state. Methods in this study can be applied in future studies using serial nasal secretion collections to evaluate important pharmacodynamic indices linked to efficacy, for example percentage of time that free drug remains above the MIC over a 24-hour period, the ratio of free drug area under the concentration-time curve (AUC) to MIC over a 24-hour period, and the ratio of maximum concentration to MIC.

A drawback of drug concentrations reported from whole tissues or secretions is the assumption that antibiotics are uniformly distributed within tissue compartments (intracellular, interstitial, intravascular). Newly developed methods have been applied to a number of respiratory studies to evaluate the distribution of antibiotics across these compartments, which may represent different sites of infections. Future studies should evaluate intracellular levels of antibiotics in sinonasal secretions, since polymorphonuclear leucocytes and macrophages have a high uptake of macrolides and tetracyclines(47, 56, 57).

This study does not evaluate the factors influencing drug penetration of antibiotics but considers them by using a prospective randomized control design. A relatively larger unexplained variation is seen in the concentration of doxycycline in the serum and the concentration of roxithromycin in the mucus. Drug penetration of antibiotics is dependent on both drug-related and host-related factors(45, 49, 50, 57). Drug-related factors include pKa, lipophilicity, protein binding, molecular weight and mode of transport(45, 57). Host-related factors include macrophage uptake, bio-inactivation from various sources including bacterial or leucocyte enzymes and elemental ions, or elimination mechanisms including lymphatic drainage and mucociliary transport(45, 49, 50, 57). Inflammation is also an important factor since it increases the amount of tissue binding, which will inhibit movement of antibiotics across the mucosa into the mucus(57). The post-antibiotic effect which quantifies the persistence of bacterial suppression after short exposure to the drug is also unknown.

## CONCLUSIONS

The concentration of doxycycline and roxithromycin in nasal mucus was less than those in the sinonasal mucosa or systemic circulation. Based on the MIC of individual bacterial species associated with CRS these were therapeutic in the tissue and serum but not in the mucus. However, effective distribution to the infection site cannot be assumed alone on predicted bacterial susceptibilities since antibiotic resistance is variable and microbes exist in complex communities that may increase their tolerance to antibiotics. A short course of antibiotic intake did not correlate with any significant difference in endoscopic assessment nor gastrointestinal symptom scores between groups except where slightly less favourable gastrointestinal scores were found in patients taking roxithromycin. Further research is required in order to determine the factors influencing drug penetration in the mucus, and more importantly whether this is clinically relevant.

## Acknowledgements

Laboratory and technical assistance were provided by the microbiology laboratory in Labtests Auckland, and the research and development team in Zenith Technology Corporation. This study was supported by a grant from the Garnett Passe and Rodney Williams Memorial Foundation.

## Transparency declarations

None to declare.

